# Schema-Grounded Multitask Instruction Fine-tuning for Joint Biomedical Named Entity Recognition and Relation Extraction in Pharmacovigilance

**DOI:** 10.64898/2026.07.30.741807

**Authors:** Hasin Rehana, Junguk Hur

## Abstract

**Motivation:** Pharmacovigilance relies on accurate extraction of structured biomedical entities and their semantic relationships from scientific literature. However, most biomedical information extraction systems address named entity recognition (NER) and relation extraction as separate tasks trained on corpus-specific architectures, limiting scalability and cross-task knowledge sharing. Recent developments in instruction-tuned Large Language Models (LLMs) offer a promising alternative through unified generative extraction, but robust schema-grounded multitask adaptation for biomedical extraction is still understudied.

**Methods:** This study proposes a unified multitask instruction-tuned LLM framework that jointly performs biomedical NER and relation extraction across three benchmark corpora to identify chemical, disease, drug entities, as well as chemical-disease relations, drug-adverse event relations, and drug-drug interactions. Two general LLMs, Llama-3.2-3B-Instruct and Qwen3-8B, were fine-tuned using Low-Rank Adaptation (LoRA) under a shared generation interface that extracts both entity pairs and their underlying relation. Zero-shot and fine-tuned configurations were evaluated across all the tasks on their respective held-out test sets.

**Results:** Parameter-efficient fine-tuning substantially improved both entity and relation extraction performance across all tasks and model families. Fine-tuned Qwen3-8B achieved the strongest overall performance with 89.42% micro-averaged entity F1 and 62.32% micro-averaged relation F1. Fine-tuned Llama-3.2-3B achieved 87.63% entity F1 and 58.42% relation F1 despite its substantially smaller parameter count, outperforming the zero-shot 8B model on both tasks. Fine-tuning also reduced structured JSON parse failures from 23.5% to 0.11%, demonstrating stable schema internalization during supervised adaptation.

**Conclusion:** Schema-grounded multitask instruction tuning with LoRA provides a robust and computationally feasible framework for unified biomedical information extraction across heterogeneous benchmark corpora. The findings further demonstrate that schema-grounded adaptation is substantially more important than model scale alone for reliable extraction of structured biomedical relations. The gap between NER and relation extraction performance motivates future research on explicit negative-relation supervision and ontology-guided relation extraction.

## 1. Introduction

Biomedical Literature contains a rapidly expanding volume of pharmacological, toxicological, and clinical evidence, making manual curation increasingly impractical at scale [1]. Automated biomedical information extraction systems, therefore, play a critical role in pharmacovigilance, adverse drug event surveillance, drug-drug interaction monitoring, and biomedical knowledge base construction. There are two core areas for biomedical information extraction: named entity recognition (NER) and relation extraction. NER focuses on identifying and classifying drug names, chemical compounds, diseases, and adverse events in free text. On the other hand, relation extraction determines whether a semantic relationship holds between a pair of identified entities [2].

Traditional biomedical Natural Language Processing (NLP) systems usually treat NER and relation extraction as separate supervised tasks trained independently on corpus-specific architectures. Encoder-based transformer models such as BioBERT [3] and PubMedBERT [4] have achieved strong performance on biomedical NER benchmarks through domain-adaptive pretraining on PubMed and PMC corpora [5]. However, relation extraction remains substantially more challenging because it requires contextual, semantic, and causal reasoning beyond local span detection [6]. Besides, maintaining separate task-specific models introduces engineering complexity and limits knowledge transfer across related biomedical extraction tasks [7]. Moreover, maintaining separate task-specific models increases engineering complexity and limits transfer across related biomedical extraction problems.

Recent generative and end-to-end biomedical information extraction approaches attempt to jointly model entity and relation prediction through structured sequence generation directly from raw text. Seq2rel reframed relation extraction as constrained sequence generation and demonstrated that entity extraction and relation prediction could be learned jointly within a unified sequence-to-sequence framework, reducing reliance on traditional pipeline architectures [8]. BioGPT demonstrated competitive biomedical generation performance on downstream extraction tasks, including 44.98% F1 on BC5CDR relation extraction and 40.76% F1 on drug-drug interaction extraction under end-to-end settings [9]. Building on these ideas, subsequent unified extraction frameworks extended generative modeling beyond relation prediction to jointly represent entities, events, and semantic relations within a common text-to-structure generation paradigm. In parallel, parameter-efficient fine-tuning (PEFT) [10] methods such as LoRA [11] and QLoRA [12] have substantially reduced the computational cost of adapting large language models to domain-specific tasks, making biomedical instruction tuning feasible under modest hardware constraints [13].

Despite these advances, unified multitask biomedical information extraction across heterogeneous corpora remains substantially more difficult than isolated pipeline prediction. Biomedical corpora differ substantially in annotation granularity, relation taxonomies, and schema structure. In pharmacovigilance-oriented corpora, relation prediction is further confounded by strong co-occurrence priors that can lead to false-positive causal predictions when entities appear in the same sentence or abstract without an explicit semantic relation [14].

This paper presents a schema-grounded multitask instruction tuning framework for joint biomedical NER and relation extraction across three benchmark corpora: BC5CDR [15], ADE V2 [16], and DDI-2013 corpora [17]. The framework casts all tasks as constrained JSON generation under a shared prompting interface while preserving corpus-specific schema constraints and relation definitions. The primary contributions of this work are:

1. A unified multitask biomedical information extraction corpus combining BC5CDR, ADE Corpus V2, and DDI-2013 under a shared schema-grounded generative framework.
2. Task-specific instruction design incorporating co-occurrence guards and relation disambiguation cues for heterogeneous biomedical relation schemas.
3. Parameter-efficient LoRA fine-tuning of Llama-3.2-3B-Instruct and Qwen3-8B under a shared structured generation interface.

## 2. Methods

### 2.1 Study Design and Task Formulation

We developed a unified multitask instruction-tuned framework for biomedical information extraction in pharmacovigilance-oriented scientific text. The framework jointly models NER and relation extraction under a single structured generation interface. Given an input passage and task-specific instructions, the model generates a constrained JSON output containing entity mentions and any pairwise relations. Three extraction tasks were integrated into a unified formulation: chemical-disease extraction, drug-adverse event extraction, and drug-drug interaction extraction. This unified formulation allows the model to share lexical and semantic knowledge across tasks while preserving task-specific label constraints through schema-aware prompting. **Figure 1** demonstrates the overall pipeline of this research.

**Figure 1.**
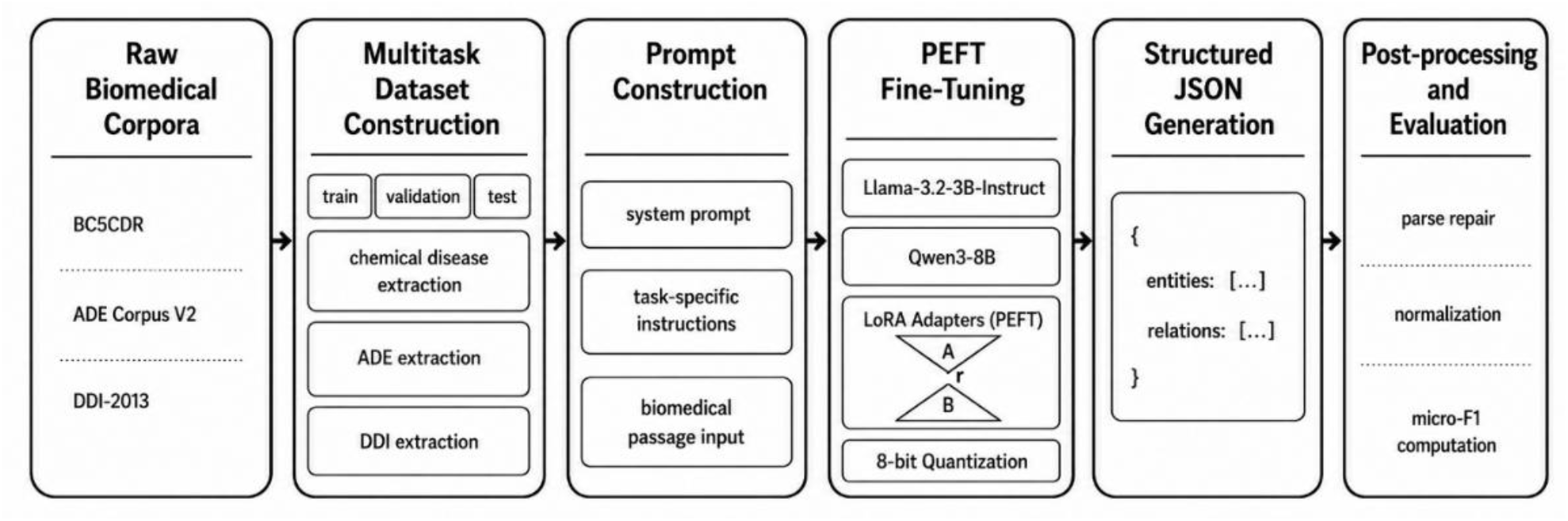
Unified multitask biomedical information extraction pipeline. The framework integrates preprocessing, prompt construction, LoRA fine-tuning, structured generation, and evaluation.

### 2.2 Dataset

The multitask dataset combined three publicly available biomedical information extraction benchmarks: BC5CDR [15], ADE V2 [16], and DDI-2013 [17] corpora. Each corpus contained both named entity and relation annotations for joint extraction training.

BC5CDR had PubMed abstracts annotated for chemical and disease entities and chemical-induced disease relations. Because BC5CDR relations were defined at the document level, relations were resolved to passage-level instances during preprocessing.

ADE V2 had drug and adverse event entities with drug-adverse event relations extracted from MEDLINE case reports. PMID level splitting was preserved to prevent sentence-level contamination. DDI-2013 contained drug entity annotations for four relation subtypes (mechanism, effect, advise, int) derived from SemEval-2013. The final multitask dataset consisted of 9,943 training examples, 1,107 validation examples, and 2,686 test examples, shown in **Table 1**.

**Table 1.** Dataset statistics for the unified multitask biomedical information extraction corpus. BC5CDR, ADE V2, and DDI-2013 were merged into a unified multitask dataset while preserving corpus-specific relation schemas. BC5CDR had a higher test ratio (33%) because its official test set equals the train set size. Changing this would break benchmark comparability. CID: Chemical-induced disease, ADE: Adverse-drug event, DDI: Drug-drug-interaction.

| Corpus | Entity Types | Relation Type | Task | Train | Val | Test |
| --- | --- | --- | --- | --- | --- | --- |
| BC5CDR | Chemical, Disease | Chemical-induced disease | NER + CID RE | 1,777 | 198 | 945 |
| ADE V2 | Drug, Adverse Event | Adverse Drug Event | NER + ADE RE | 3,252 | 361 | 657 |
| DDI-2013 | Drug | Drug-drug interaction | NER + DDI RE | 4,999 | 555 | 1,084 |
| Total |  |  |  | 9,943 | 1,107 | 2,686 |

### 2.3 Task Formulation

All biomedical extraction tasks were formulated as conditional structured text generation. Given a biomedical passage and task instruction, the model generated a single JSON object that contains predicted entities and relations. Each entity record included text span, entity type and character offsets. For chemical-induced disease relation extraction, the instruction explicitly states that chemical-disease co-occurrence is insufficient evidence for a chemical-induced disease relation; a direct causal or inductive link must be present. For adverse drug events, the instruction defines adverse drug events to exclude therapeutic effects. For drug-drug interaction, the instruction provides taxonomic definitions for all four relation subtypes. This schema design ensures the model is never required to produce null-valued cross-task fields, which is a key distinction from naive multitask training targets.

### 2.4 Prompting schema and structured JSON output constraint

All three tasks share a common system prompt instructing the model to output raw JSON only, with no markdown or explanatory text. Each prompt receives an additional task-specific instruction defining entity types, relation types, and disambiguation cues relevant to that domain. Chemical-induced disease identification prompts emphasized causal evidence requirements for chemical-induced disease relations, whereas drug-drug interaction identification prompts included subtype-specific semantic cues for interaction classification.

Structured generation constraints were designed to stabilize decoding behavior while preserving flexibility across heterogeneous biomedical relation schemas. **Figure 2** illustrates the shared prompting structure and task-conditioned JSON generation framework used across the three biomedical extraction tasks.

**Figure 2.**
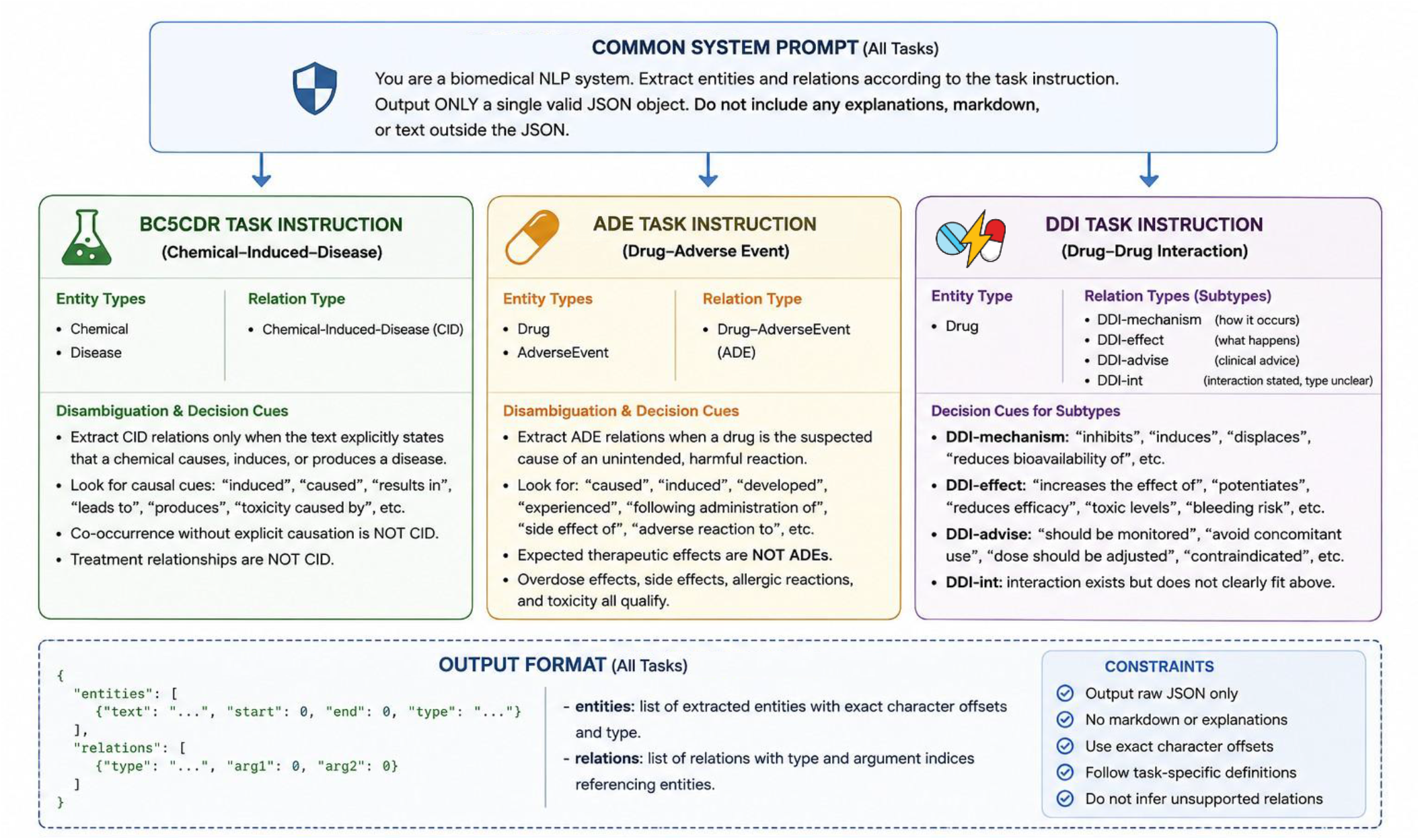
Task-specific prompting schema and structured JSON output format. A unified generative framework was used across BC5CDR, ADE V2, and DDI-2013 through task-conditioned prompting and constrained JSON decoding.

### 2.5 Model Architecture and Parameter-Efficient Fine-Tuning

We evaluated two instruction-tuned LLMs: Llama-3.2-3B-Instruct [18] and Qwen3-8B [19]. Both models were adapted using Low-Rank Adaptation (LoRA) with 8-bit quantization. Details of the hyperparameters are listed in **Table 2**. Adaptation layers were applied to seven transformer target modules. The Llama-3.2-3B configuration contained 48.6M trainable parameters, corresponding to approximately 1.49% of total model parameters.

**Table 2.** Training hyperparameters for LoRA fine-tuning. Parameter-efficient adaptation was performed using 8-bit quantization and shared optimization settings.

| Parameter | Value |
| --- | --- |
| Base model | Llama-3.2-3B-Instruct |
| Quantization | 8-bit BitsAndBytes (fp16 compute) |
| LoRA r / alpha | 32 / 64 |
| Learning rate | 1e-4 |
| LR scheduler | Cosine with 5% warmup |
| Epochs | 5 |
| Per-device batch size | 2 |
| Gradient accumulation | 4 |
| Max sequence length | 2,048 |
| Optimizer | paged_adamw_8bit |

Training employed cosine learning-rate scheduling with checkpoint selection based on validation loss. An effective batch size was achieved through gradient accumulation across multi-GPU training. Llama-3.2-3B fine-tuning was conducted on 8× NVIDIA V100 16GB GPUs, while Qwen3-8B fine-tuning used 3× NVIDIA A800 40GB GPUs.

### 2.6 Inference pipeline and parsing

Inference was conducted through a sharded parallel evaluation pipeline. The test split was stride-interleaved into evaluation shards to support distributed decoding across multiple GPUs. Predictions from all shards were subsequently merged into a single evaluation file prior to scoring. As generative extraction systems may produce malformed structured outputs, we incorporated an explicit post-generation repair stage. A dedicated parser repair script corrected recoverable JSON formatting errors, primarily missing-comma failures. The repair stage was intentionally conservative and applied only to syntactically recoverable outputs. Structured generation stability was evaluated through parse-failure analysis.

### 2.7 Evaluation metrics

All systems were evaluated using a case-insensitive, schema-tolerant scorer that is applied uniformly across the three tasks. Entity matches required exact text equality after case normalization and must agree on the predicted entity type. Case-insensitive matching was used to reduce spurious penalization arising from capitalization inconsistencies in generative outputs. Relation matches require a correct relation type and alignment of subject and object entities to their corresponding predicted entities. The evaluation pipeline additionally incorporated a fallback matching strategy that normalizes relation labels and tolerates minor variations in output format. Micro-averaged precision, recall, and F1 scores were computed across the entire multitask test set by aggregating entity and relation counts across all three biomedical extraction tasks.

## 3. Results

All reported metrics (**Table 3**) are computed over a held-out test partition of 2,686 examples derived from the combined BC5CDR, ADE V2, and DDI-2013 corpora. Across both model families, parameter-efficient fine-tuning substantially improved joint biomedical entity and relation extraction performance relative to zero-shot prompting. The strongest overall results were obtained by the fine-tuned Qwen3-8B model, which achieved 89.42% micro-averaged entity F1 and 62.32% micro-averaged relation F1.

**Table 3.** NER and relation extraction performance across BC5CDR, ADE V2, and DDI-2013 for zero-shot and fine-tuned configurations of Llama-3.2-3B and Qwen3-8B on the held-out multitask test set (n = 2,686).

| Dataset | Task | NER |  |  | RE |  |  |
| --- | --- | --- | --- | --- | --- | --- | --- |
|  | Model | Precision (%) | Recall (%) | F1 Score (%) | Precision (%) | Recall (%) | F1 Score (%) |
| BC5CDR | Llama-3B ZS | 41.33 | 37.16 | 39.14 | 39.39 | 6.94 | 11.81 |
|  | Llama-3B FT | 85.42 | 83.81 | 84.61 | 44.58 | 49.32 | 46.83 |
|  | Qwen3-8B ZS | 56.01 | 59.58 | 57.74 | 35.00 | 23.74 | 28.29 |
|  | Qwen3-8B FT | **86.46** | **86.44** | **86.45** | **48.1** | **48.78** | **48.44** |
| ADE V2 | Llama-3B ZS | 65.48 | 70.28 | 67.79 | 62.44 | 38.48 | 47.61 |
|  | Llama-3B FT | 88.01 | 84.00 | 85.96 | 78.98 | 72.49 | 75.60 |
|  | Qwen3-8B ZS | 82.75 | 76.18 | 79.33 | 72.08 | 58.17 | 64.38 |
|  | Qwen3-8B FT | **88.49** | **87.02** | **87.74** | **79.87** | **77.77** | **78.81** |
| DDI-2013 | Llama-3B ZS | 79.11 | 61.89 | 69.45 | 13.62 | 37.19 | 19.94 |
|  | Llama-3B FT | 94.49 | 93.57 | 94.02 | 60.02 | 66.05 | 62.89 |
|  | Qwen3-8B ZS | 91.88 | 80.37 | 85.74 | 39.81 | 17.95 | 24.74 |
|  | Qwen3-8B FT | **95.40** | **96.07** | **95.73** | **71.56** | **69.62** | **70.58** |
| Micro Average | Llama-3B ZS | 56.22 | 50.36 | 53.13 | 24.50 | 23.37 | 23.92 |
|  | Llama-3B FT | 88.57 | 86.72 | 87.63 | 56.95 | 59.97 | 58.42 |
|  | Qwen3-8B ZS | 69.89 | 68.67 | 69.27 | 48.37 | 31.68 | 38.29 |
|  | Qwen3-8B FT | **89.47** | **89.38** | **89.42** | **62.56** | **62.09** | **62.32** |

The fine-tuned Llama-3.2-3B model achieved 87.63% entity F1 and 58.42% relation F1 despite its substantially smaller parameter count. Zero-shot performance was consistently lower for both architectures. Llama-3.2-3B zero-shot achieved 53.13 entity F1 and 23.92 relation F1, while Qwen3-8B zero-shot achieved 69.27 entity F1 and 38.29 relation F1. These results indicate that instruction alone is insufficient for reliable structured biomedical extraction under the proposed unified schema.

Fine-tuning produced substantial improvements for the 3B model family. Relative to Llama-3.2-3B zero-shot prompting, LoRA adaptation improved entity F1 from 53.13% to 87.63% and relation F1 from 23.92% to 58.42%. Notably, the fine-tuned 3B model also outperformed the zero-shot 8B model on both entity and relation extraction despite using fewer parameters, suggesting that task-specific structured adaptation was more influential than model scale alone in this setting.

Figure 3 summarizes both the performance gains obtained through LoRA adaptation and the residual relation prediction biases observed across datasets. Fine-tuning substantially improved both entity and relation extraction across all tasks. However, relation prediction behavior remained dataset-dependent. BC5CDR retained a mild false-positive bias after fine-tuning, consistent with persistent co-occurrence-driven errors, whereas Llama-3.2-3B zero-shot exhibited substantial over-prediction behavior on drug-drug interaction extraction.

Task-level results exhibited consistent trends across model variants. NER performance was generally stronger than relation extraction, particularly in the zero-shot setting. Chemical-induced disease identification was consistently the most difficult relation extraction domain across all evaluated models. Although fine-tuned models achieved strong NER performance on BC5CDR, relation extraction performance remained comparatively weaker, likely because chemical-disease co-occurrence alone is often insufficient evidence for a causal chemical-induced disease relation.

**Figure 3.**
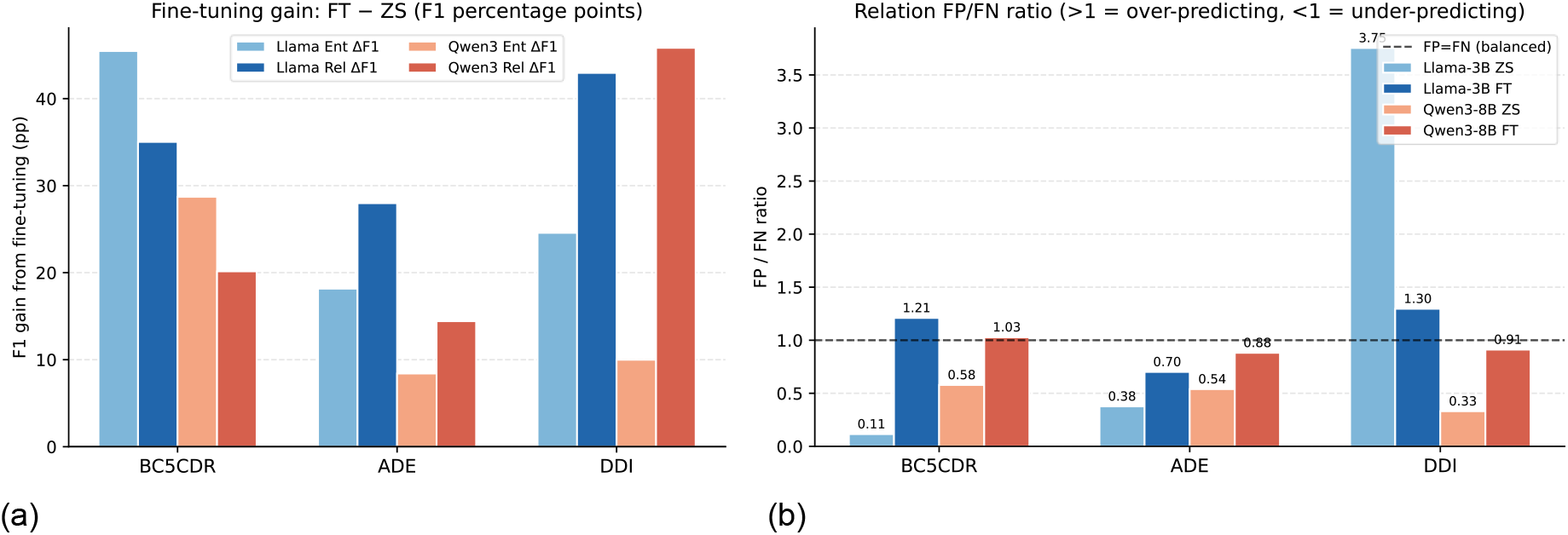
Fine-tuning gains and relation prediction behavior across biomedical extraction tasks. (a) F1 improvements after LoRA fine-tuning relative to zero-shot prompting. (b) False-positive to false-negative ratio for relation extraction across datasets and model configurations. Ent: Entity, Rel: Relation, ZS: Zero Shot, FT: Fine-tuned.

Adverse drug event relation extraction was substantially easier under the current formulation. Fine-tuned Llama-3.2-3B achieved 75.60% F1 for adverse drug event, and fine-tuned Qwen3-8B achieved 78.81% relation F1. Adverse drug event relation extraction also yielded the strongest zero-shot relation performance among the three relation extraction, with Qwen3-8B zero-shot reaching 64.38% relation F1.

NER performance on DDI-2013 dataset was high for fine-tuned models, reaching 94.02 entity F1 for Llama-3.2-3B FT and 95.73 entity F1 for Qwen3-8B FT. However, zero-shot drug-drug interaction extraction remained weak despite relatively strong entity detection, indicating that interaction subtype classification was more difficult than mention identification alone. Fine-tuning produced particularly large improvements for drug-drug interaction extraction, with fine-tuned Qwen3-8B achieving the strongest overall drug-drug interaction identification performance.

Fine-tuning also dramatically improved structured decoding reliability, reducing the parse failure (**Table 4**), which is a critical prerequisite for practical generative biomedical IE systems. Llama-3.2-3B zero-shot produced 630 malformed outputs (23.5%), whereas the fine-tuned model produced only 3 unrecoverable parse failures (0.11%). Qwen3-8B exhibited near-zero parse failures in both zero-shot and fine-tuned settings. Qwen3-8B demonstrates substantially more robust zero-shot structured generation (2 failures; 0.07%). The near-zero post-fine-tuning failure rates for both families indicate that the learned output schema is stably internalized and generalizes reliably to unseen test instances.

**Table 4.** Structured JSON parse failures on the test set (n = 2,686). Fine-tuning dramatically improved structured decoding reliability and reduced malformed outputs.

| Model | Parse Failures |
| --- | --- |
| Llama-3.2-3B ZS | 630 |
| Llama-3.2-3B FT | 3 |
| Qwen3-8B ZS | 2 |
| Qwen3-8B FT | 1 |

## 4. Discussion

This study demonstrates that multitask instruction tuning with PEFT provides a robust framework for unified biomedical information extraction across heterogeneous corpora. Across all tasks, fine-tuned models substantially outperformed their zero-shot counterparts for both NER and relation extraction.

The fine-tuned Llama-3.2-3B model outperformed the zero-shot Qwen3-8B model on both entity and relation extraction despite having substantially fewer parameters. This finding suggests that task-specific adaptation and structured output supervision contribute more directly to the reliability of biomedical extraction than scaling model size alone under zero-shot prompting conditions.

RE remained substantially more difficult than NER across all settings. This discrepancy was particularly pronounced in the zero-shot setting, where models frequently identified biomedical entities correctly while failing to infer accurate relation semantics. One likely explanation is that biomedical NER is primarily a local span-identification problem, whereas relation extraction requires deep reasoning about semantic, causal, and contextual dependencies among entities. In pharmacovigilance literature, entity mentions are often lexically explicit, but their relationships may be implicit, speculative, negated, or confounded by simple co-occurrence.

Chemical-induced disease relation extraction remained the most challenging task after fine-tuning. Models frequently predicted false positive chemical-induced disease relations happed due to chemical disease co-occurrences in the absence of direct causal evidence. This pattern indicates that co-occurrence over-prediction constitutes a dominant error mode in chemical-induced disease extraction.

Drug-drug interaction subtype prediction exhibited a distinct failure pattern under zero-shot prompting. Although zero-shot models often recognized interacting drug mentions correctly, subtype assignment remained weak, suggesting that semantic distinctions among mechanism, effect, advise, and int relations require stronger supervised grounding than entity identification alone.

Fine-tuning also substantially improved the reliability of structured output. Zero-shot generation frequently produced malformed JSON outputs, especially with the smaller base model, whereas fine-tuned models produced stable, schema-consistent predictions. This finding is operationally important because downstream biomedical pipelines depend not only on semantic accuracy but also on reliable structured outputs for automated postprocessing and scalable integration.

Overall, the findings support multitask instruction tuning with LoRA as a practical and scalable strategy for unified biomedical NER and relation extraction. While relation reasoning remains challenging, compact instruction-tuned models can achieve strong multitask extraction performance with relatively modest computational resources.

The proposed framework has several practical limitations. The study did not include single-task vs. multitask ablations, preventing direct quantification of cross-task transfer effects within the unified framework. The experiments focused on two instruction-tuned causal LLM families and did not evaluate broader architectural alternatives. In addition, the evaluation focused on exact-match metrics without any human qualitative assessment of the quality of the generated assessment.

Future work should focus on reducing co-occurrence bias through hard-negative training and integrating ontology knowledge base into the fine-tuning pipeline.

## 5. Conclusion

This research presents a unified schema-grounded multitask instruction fine-tuning framework for joint biomedical NER and relation extraction across BC5CDR, ADE V2, and DDI-2013 corpora. Parameter-efficient LoRA adaptation substantially improved both extraction accuracy and structured decoding reliability across all evaluated tasks and model families. The fine-tuned 3B model surpassed the zero-shot 8B model on both tasks, demonstrating the importance of supervised, schema-grounded adaptation over model scale alone under zero-shot prompting conditions.

RE remains substantially more difficult than NER, particularly for chemical-induced disease prediction where co-occurrence bias persisted despite improved NER performance. The study also emphasizes the importance of structured generation reliability in biomedical NLP pipelines. Fine-tuning improved not only extraction accuracy but also schema consistency, substantially reducing the number of malformed structured outputs.

Future work will prioritize hard-negative training to address co-occurrence bias at the supervision level. Preliminary experiments indicate that post hoc ontology filtering alone does not resolve this issue. Broader evaluation across additional corpora and model families is also warranted.

## 6. Acknowledgement

The study was supported by the U.S. National Institute of Allergy and Infectious Disease (U24AI171008 to J.H.).

